# Multiple Isoforms of the Activin-like receptor Baboon differentially regulate proliferation and conversion behaviors of neuroblasts and neuroepithelial cells in the *Drosophila* larval brain

**DOI:** 10.1101/2024.03.05.583454

**Authors:** Gyunghee G. Lee, Aidan J. Peterson, Myung-Jun Kim, Michael B. O’Connor, Jae H. Park

## Abstract

In *Drosophila* coordinated proliferation of two neural stem cells, neuroblasts (NB) and neuroepithelial (NE) cells, is pivotal for proper larval brain growth that ultimately determines the final size and performance of an adult brain. The larval brain growth displays two phases based on behaviors of NB and NEs: the first one in early larval stages, influenced by nutritional status and the second one in the last larval stage, promoted by ecdysone signaling after critical weight checkpoint. Mutations of the *baboon* (*babo*) gene that produces three isoforms (BaboA-C), all acting as type-I receptors of Activin-type transforming growth factor β (TGF-β) signaling, cause a small brain phenotype due to severely reduced proliferation of the neural stem cells. In this study we show that loss of *babo* function severely affects proliferation of NBs and NEs as well as conversion of NEs from both phases. By analyzing *babo*-null and newly generated isoform-specific mutants by CRISPR mutagenesis as well as isoform-specific RNAi knockdowns in a cell- and stage-specific manner, we further demonstrate differential contributions of the isoforms for these cellular events with BaboA playing the major role. Our data show that stage-specific expression of EcR-B1 in the brain is also regulated primarily by BaboA along with function of the other isoforms. Blocking EcR function in both neural stem cells results in a small brain phenotype that is more severe than *baboA*-knockdown alone. In summary, our study proposes that the Babo-mediated signaling promotes proper behaviors of the neural stem cells in both phases and achieves this by acting upstream of EcR-B1 expression in the second phase.

**Author Summary:** Evolutionarily conserved TGF-β signaling pathway is widely utilized as a regulator of diverse processes of brain development in both vertebrates and invertebrates. A key element in *Drosophila* Activin type TGF-β signaling pathway is the type-I receptor Babo. The *babo* gene produces three isoforms, each with a unique ligand preference. Our study uncovers that Babo-mediated signaling promotes proper proliferation of NBs and NEs and conversion of NEs, together responsible for the magnitude of larval brain size growth. Three isoforms act individually or together to regulate these cellular events in coordination with developmental status. Our findings emphasize that Babo-mediated signaling is a crucial regulator of postembryonic neurogenesis that generates 90% of neuronal population for the adult brain.

## Introduction

In mammals, the brain grows not only during prenatal development but also after birth. For example, a human brain that weighs less than a pound at birth gains about two more pounds by the early teens [1]. The postnatal brain growth highly influences the overall performance of the adult brain. In the fruit fly *Drosophila melanogaster,* postembryonic brain growth is also crucial for establishing the adult brain.

When a newly hatched larva increases its body weight about 1000-fold by the end of larval development, its brain also undergoes an incredible size increase due to a rapid accumulation of many immature neurons produced by postembryonic neurogenesis that provide nearly 90% of the neuronal population of the adult brain [2]. For proper shaping and functioning of the adult brain, postembryonic neurogenesis is precisely regulated in time and space by both locally and globally acting factors. The *Drosophila* brain has proven to be an excellent system to explore underlying mechanisms of post-embryonic neurogenesis [2–7].

The *Drosophila* larval brain lobe is divided into two compartments, the central brain (CB) and the optic lobe (OL). The same set of approximately 100 neuroblasts (NBs) is responsible for both embryonic and postembryonic neurogenesis in the CB [8–9]. Toward the end of embryonic development, the NBs become dormant and then resume mitotic activities at late first-instar (L1) stage and some of them continue to proliferate until the end of the prepupal stage [2,10]. The only exception to this pattern is mushroom body (MB) NBs that divide without dormancy [11].

Larval feeding on protein-rich food is known to promote the fat body to release an unknown factor, which triggers the dormant NBs to resume their mitotic activity [12–15]. During this phase, NBs can alter the cell cycle status in a nutrient-dependent manner, showing their plasticity in response to dietary conditions [16]. Under normal growth conditions, the critical weight (CW) checkpoint is about 8-10 h after ecdysis into the third-instar stage (L3) when the larvae attain about 0.75 mg in weight [17–19].

After the CW checkpoint, NBs divide faster, marking their entry into the second growth phase. Starvation after CW checkpoint accelerates the time to pupariation, but NBs maintain their second phase proliferation rate/mode, indicating that their proliferation becomes insensitive to nutritional conditions [2,10,20]. Ecdysone/Ecdysone receptor (EcR) signaling is suggested to play a key role in promoting NBs passage into the second growth phase [21–22].

The growth of the larval OL differs from that of the CB. The OL primordium of a newly hatched larva contains only NEs that have migrated from the posterior procephalic ectoderm during embryogenesis and is organized into two distinct layers, the outer (OPC) and inner proliferation centers (IPC) [23]. The NEs also behave differently in two distinct phases: symmetric divisions to expand the NE population in first phase, followed by their gradual transformation into NBs and other precursor neuronal populations in the second phase [2]. Like NBs in the CB, the proliferation of NEs in the first phase is influenced by dietary conditions. In the OPC, the second phase is hallmarked by the gradual conversion of NEs into medulla NBs medially in a medial-to-lateral direction as “proneuronal waves” and lamina precursor cells (LPC) laterally [24]. The timing of OPC NE conversion is known to be influenced by various signals including the JAK/STAT, Hippo/Yorkie, Notch, and EGFR signaling pathways [25–31]. It has also been suggested that ecdysone/EcR signaling orchestrates unfolding of second phase events [22,32,33]; however, little is known about how these various signals are coordinated by ecdysone/EcR signaling.

The *babo* gene encodes the only type-I receptor of the Activin-branch of the transforming growth factor-β (TGF-β) signaling pathway and plays a key role in regulating EcR-B1 expression in the larval CNS [18]. High levels of EcR-B1 expression are found in diverse cell types in the CNS at late L3 stage; however, such EcR-B1 expression is mostly abolished in *babo*-null (pan-*babo*) mutants lacking functions of all isoforms, suggesting that Babo-mediated signaling regulates stage-specific EcR-B1 expression in the CNS. In line with this, pan-*babo* mutant larvae exhibit a small brain phenotype caused by greatly reduced postembryonic neurogenesis [34,35]. The *babo* locus produces three isoforms, BaboA, BaboB, and BaboC, differing only in their ligand binding domains. The three ligands in the Activin branch of the TGF-β family, Myoglianin (Myo), Activin-β (Actβ), and Dawdle (Daw), are thought to interact preferentially with BaboA, BaboB, and BaboC, respectively [36–38]. According to the conventional model, ligand-bound Babo forms a heteromeric complex with one, or a combination, of the type-II receptors, Punt and Wishful-thinking [38]. In the canonical TGF-β signaling, ligand-bound receptors phosphorylate dSmad2, which enters the nucleus with the co-Smad Medea to regulate gene expression. In this study, we investigated roles of pan-Babo and individual isoforms in regulating brain growth and EcR-B1 expression in the brain of L3 stage larva.

## Results

### Defective brain growth of pan-*babo* mutants

Previous studies have shown that various *babo* mutants are defective in many aspects of larval and pupal growth and development including small brains and small imaginal discs [34]. Particularly, a *trans*-heterozygous combination of loss-of-function alleles, *babo^Df^* and *babo^Fd4^*, exhibits the most severe mutant phenotypes [39]. In this study, we revisited the small-brain phenotype of *babo^Df/Fd4^*(pan-*babo*) mutants to comprehend this gene’s function in larval brain growth. At first glance, pan-*babo* mutant brains, immuno-stained with anti-Dlg to label cell types involved in postembryonic neurogenesis, appear substantially smaller than control brains at the wandering third-instar larval stage (WL3) just before puparium formation (Fig 1A and 1B). We further measured brain lobe areas of the two genotypes at 96 h and 120 h after egg laying (AEL), which represent early-to-mid and late L3 stages for control, respectively. For this analysis, we used frontal midsection confocal images of individual brain lobes because this section shows the largest area as well as the well-defined organization of the OL structures. The pan-*babo* mutant brains exhibit about 47% size reduction at 96 h AEL and about 55% at 120 h AEL (Fig 1C-1H, 1Ci-1Fi). We reasoned that a more severe reduction at 120 h AEL reflects rapid growth of the control brain but substantially retarded growth of the mutant brain during L3 development. Marked reductions are found in both the CB and OL regions of mutant brains, signifying that retarded growth of both regions contributes to the small brain phenotype (Fig 1A-1F).

**Fig 1.**
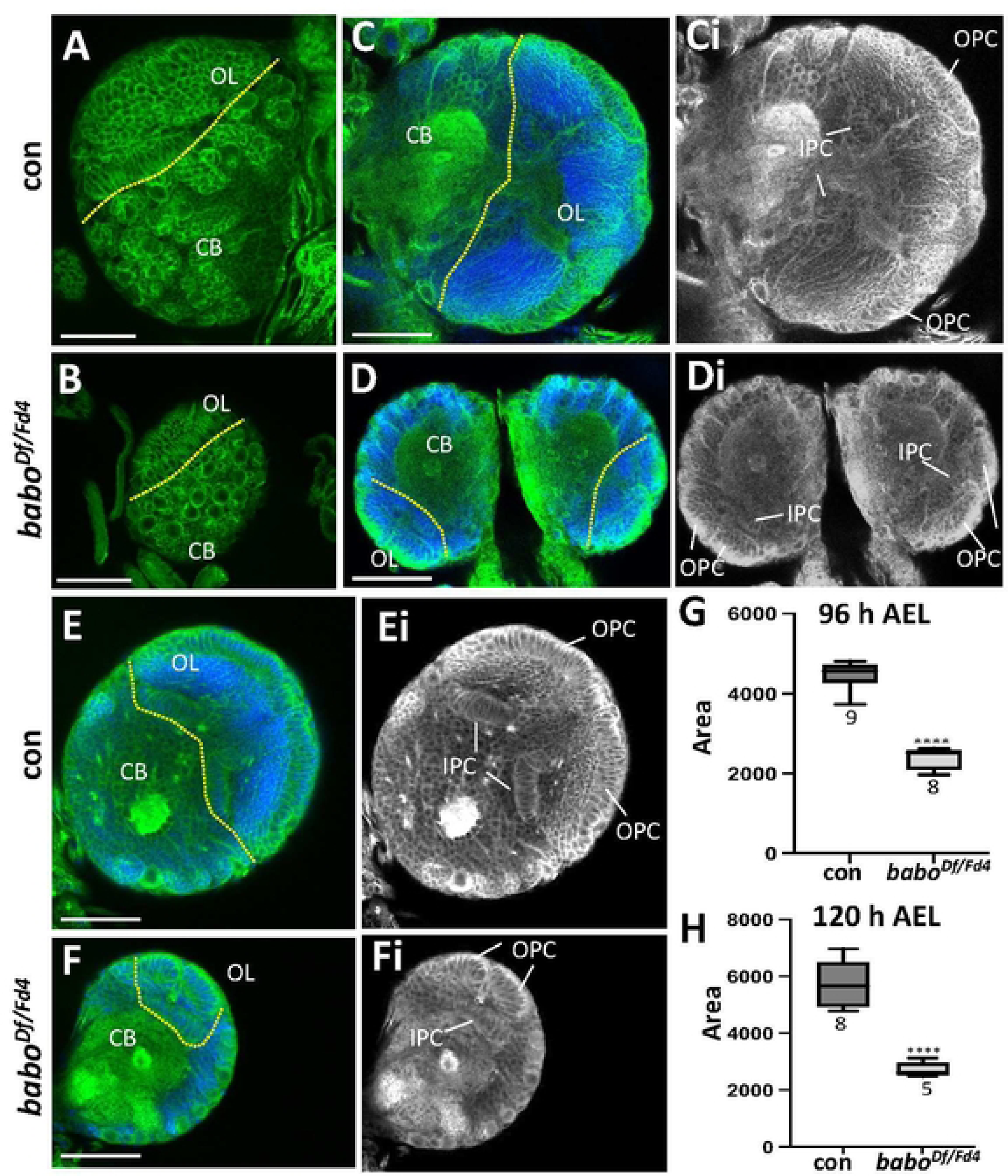
Defective brain growth of pan-*babo* (*babo^Df/Fd4^*) mutants. The dotted lines mark the border between the optic lobe (OL) and central brain (CB). Either *w^1118^* or *babo^Df/+^* is used as wildtype or genetic control, respectively. **(A,B)** Dorsal views of Dlg-labeled *babo^Df/Fd4^*and *w^1118^* (con) brains dissected at WL3 stage. **(C,D)** At 120 h AEL. Frontal mid-sections of *babo^Df/Fd4^* and *babo^Df/+^* (con) brain lobes showing Dlg and DAPI staining. **(Ci,Di)** The same images as in (C-D) showing Dlg signals alone (grey). **(E,F)** At 96 h AEL. Frontal mid-sections of *babo^Df/Fd4^* and *babo^Df/+^* (con) brain lobes co-labeled for Dlg and DAPI. **(Ei-Fi)** The same images as in (E-F) showing Dlg signals only. **(G,H)** Box plots showing mean areas (± sd) of *babo^Df/Fd4^* and con (*babo^Df/+^*) brain lobes at 96 h (G) and 120 h AEL (H). Asterisks indicate statistical significance between con and mutant. Sample sizes are indicated under each box. Scale bars in A-F: 50 μm.

### Brain growth of Individual isoform mutations

The *babo* locus produces three isoforms via alternative splicing of mutually exclusive 4^th^ exons that encode distinct ligand-binding domains (Fig 2A). To elucidate functions of individual isoforms involved in regulating brain growth, we investigated the brain phenotypes of two independent deletion mutations for each isoform: one carrying a premature stop codon and the other resulting in either a frameshift or in-frame small deletion (Fig 2B and S1). These mutations, collectively referred to as ‘indels, were assessed for brain growth defects at 120 h AEL. Intriguingly, *A-indel* and *C-indel* mutants show smaller brains than control, while *B-indel* mutant brains are unaffected in their appearance and size (Fig 2C-2F). We compared mean areas of three different brain regions (entire lobe, CB, and OL) between control and isoform mutants. *A-indel* mutations cause about a 40% reduction in the entire brain lobe area resulting from considerable reduction in both the CB and OL regions, while *C-indel* mutants show about 18% reduction mostly in the OL (Fig 2G). *B-indel* mutants do not show any significant size difference in both CB and OL regions (Fig 2G). The two independent mutations of each isoform gave very similar results, indicating that they can be used interchangeably. Taken together, these results suggest that A and C isoforms regulate different aspects of larval brain growth, while B isoform function is dispensable for brain size growth when functions of other isoforms are present.

**Fig 2.**
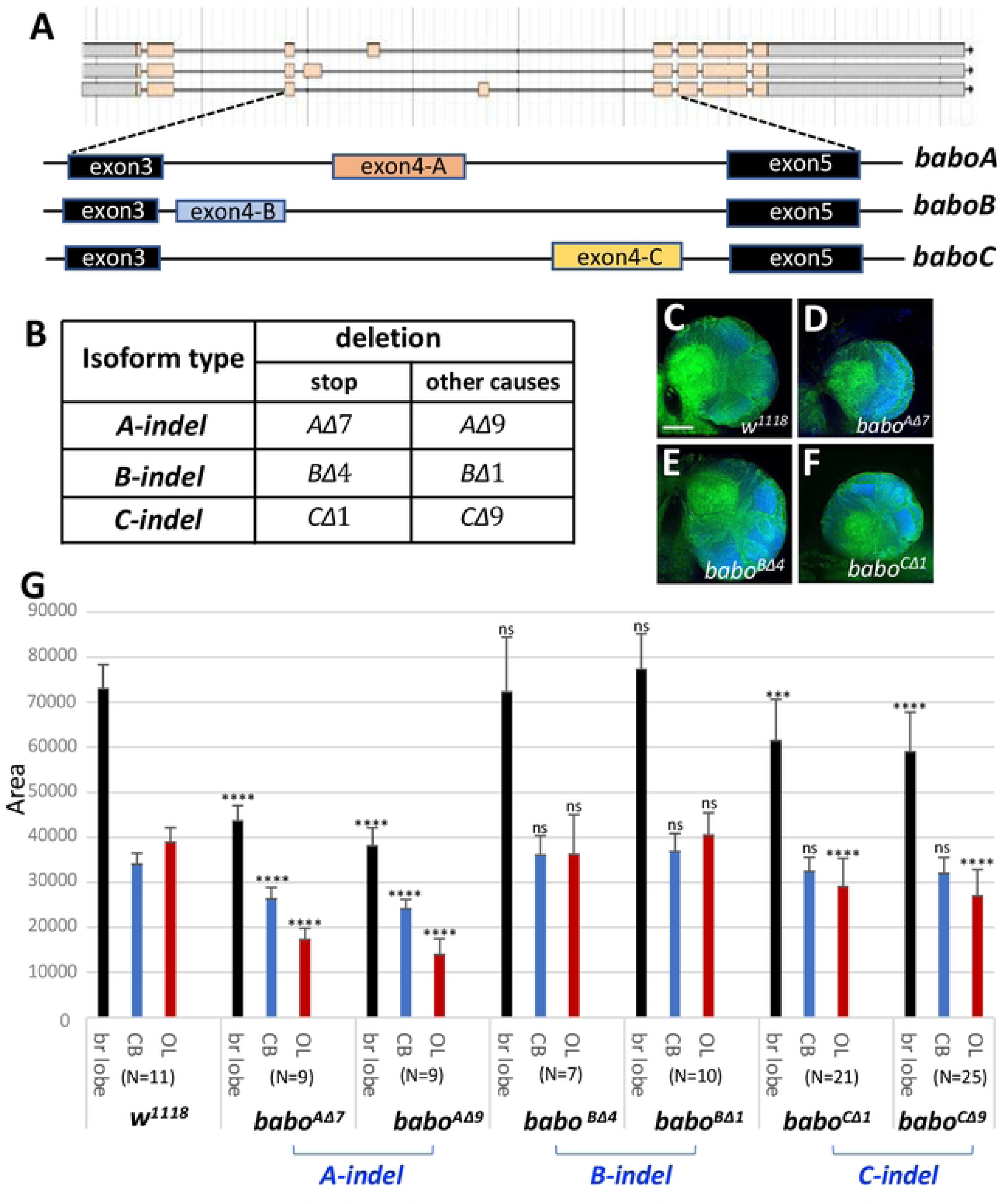
Brain size analysis of isoform-specific mutants at 120 h AEL. **(A)** Transcripts of the three *babo* isoforms resulting from alternative splicing of the 4^th^ exons (exon4-A, -B, and -C). **(B)** Two different mutation types produced by CRISPR/Cas9-mediated deletions in the isoform-specific 4^th^ exons. One type carrying stop codons, and the other causing other lesions (with or without frameshift). Both types are referred to as *indel* mutations. Details are presented in S1 Fig. **(C-F)** Frontal mid-section brain lobe images of indicated genotypes in homozygosity. The brains were dissected at 120 h and processed for anti-Dlg (green) and DAPI (blue) staining. **(G)** A bar graph showing mean areas (± sd) of three regions, whole brain lobe (black), central brain (CB, blue), and optic lobe (OL, red), of indicated genotypes. Statistically significant difference between control and mutant ones is indicated above the bars. (N, sample numbers). Scale bars in A: 50 μm.

### Brain growth of *A-indel* and *C-indel* mutants

Under normal growth conditions, control larvae reach the WL3 stage at 120 h AEL and the pupal stage at 144 h AEL (Fig 3A). However, we found that *C-indel* homozygotes develop more slowly and are still wandering or just entering the early prepupal stage at 144 h AEL, suggesting that they are about 12-24 h delayed in their development (Fig 3B). At 96 h AEL, most control larvae move away from the yeast paste and burrow into the apple juice-agar medium (Fig 3C). In contrast, *C-indel* mutants at the equivalent time also stay away from the yeast paste but are mostly crawling on the surface of the medium (Fig 3D). We speculate that the altered behavior of *C-indel* mutants might be associated with delayed larval development. To see if small *C-indel* mutant brains are the result of slow larval development, we analyzed regional areas of the *C-indel* larval brains at 120 h and 144 h AEL and compared each with control larval ones at 120 h AEL. We found that at 144hr AEL *C-indel* brain are not significantly different in size from control 120 h AEL larvae (Fig 3E). These results demonstrate that the small brain size of *C-indel* mutants at 120h AEL is the result of the delayed larval development.

**Fig 3.**
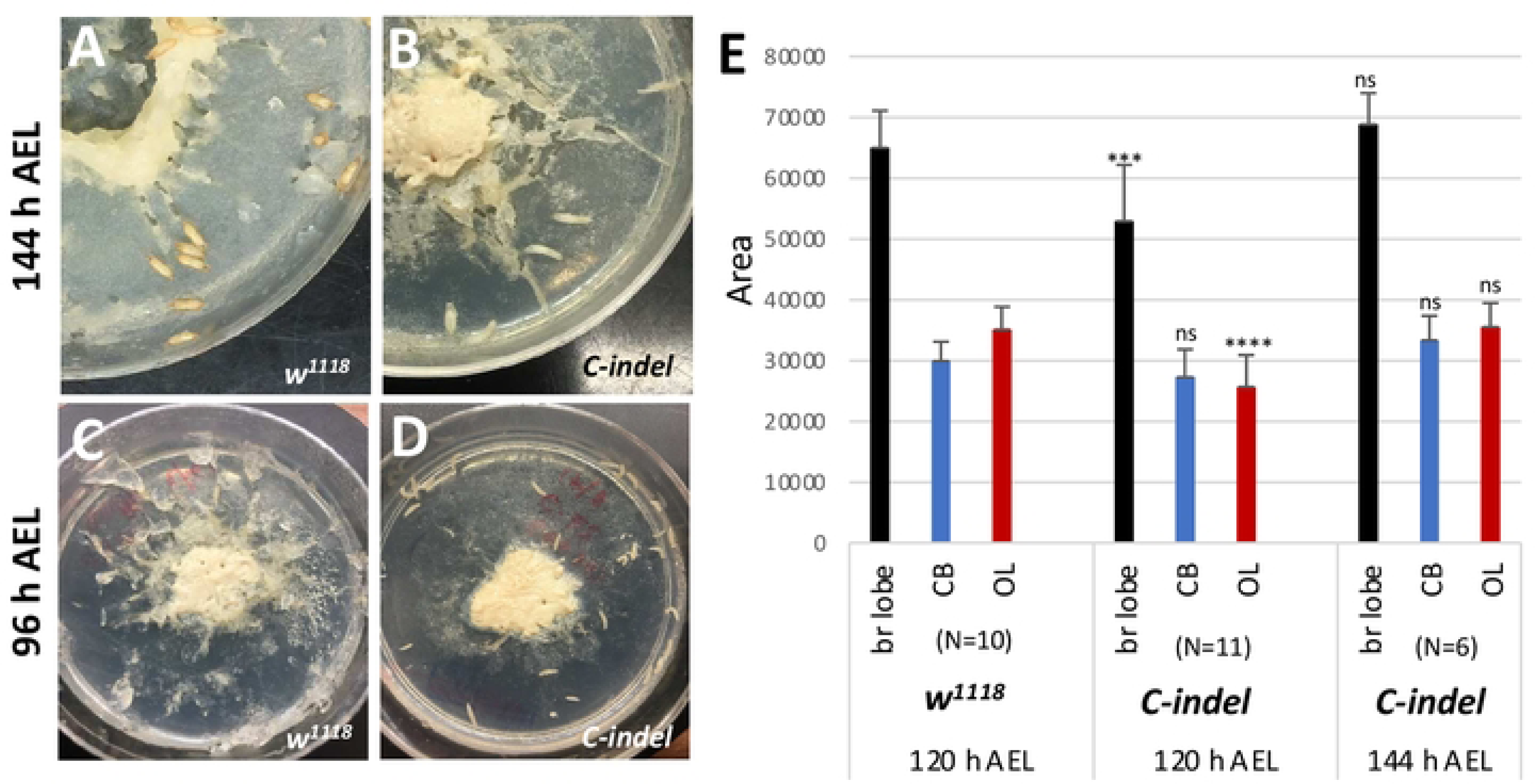
Delayed larval and brain growth of *C-indel* mutant larvae. **(A,B)** *w^1118^* and *C-indel* (*babo^CΔ1^*) mutants at 144 h AEL (n=35/plate). *C-indel* mutants displayed delayed puparium formation. **(C,D)** *w^1118^*and *C-indel* mutants at 96 h AEL. While most of *w^1118^* larvae burrowed into the agar medium, most *C-indel* mutants crawled around on the surface. **(E)** A bar graph showing mean areas (± sd) of brain lobe (black), CB (blue), and OL (red) of *w^1118^* at 120 h AEL and *C-indel* mutant larvae at 120 h and at 144 h AEL. Statistically significant differences between control and mutant are indicated above the bars. (N, sample numbers).

In contrast to *C-indel* mutants, *A-indel* mutants show normal development timing to puparium formation, but still show the small brain phenotype even at the pharate stage, strongly suggesting that BaboA function is critical for normal brain growth (Fig 4A and 4B). Defective growth is also seen in the thoracic ganglia of the ventral nerve cord (VNC) of the pan-*babo* and *A-indel* mutants (Fig 4C-4E). This is not surprising since postembryonic neurogenesis also occurs actively in the thoracic ganglia. Besides, we note that the *A-indel* mutants exhibit a less severe brain growth defect compared to pan-*babo* mutants at 120 h AEL (Fig 4C-4E). The pan-*babo* mutants delay larval development; however, their brains do not show a significant size increase even at 144 h AEL as *C-indel* mutants do and are still smaller than *A-indel* mutant larva at 120 h AEL (Fig 4F). This raises the possibility that either other isoforms act redundantly with A isoform in certain aspects of brain growth or both C and B isoforms together promote brain growth to a limited degree independently of A isoform.

**Fig 4.**
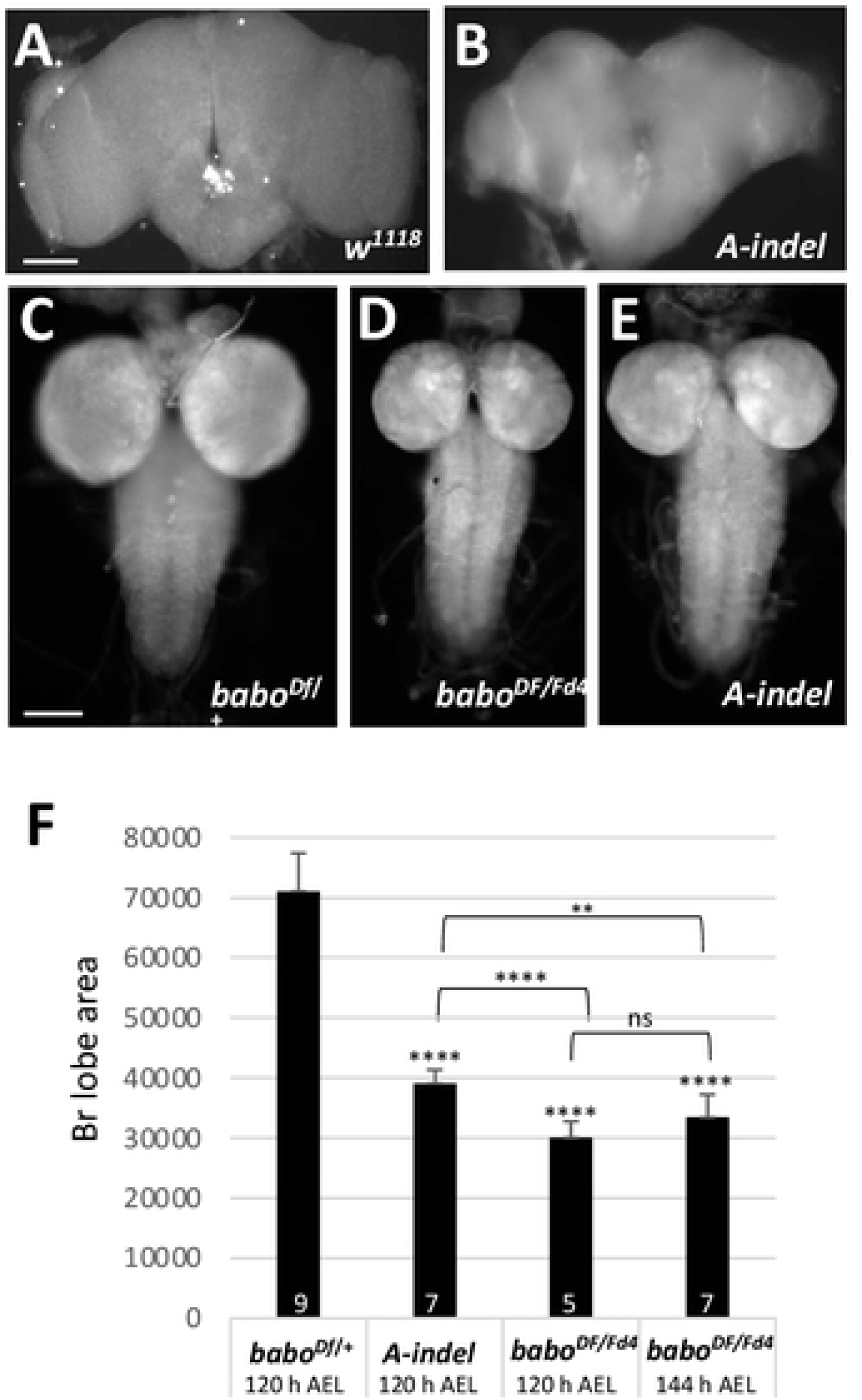
Defective brain growth of *A-indel* mutants. **(A-E)** Pharate brains of *w^1118^* (n=5) and *A-indel* (*babo^Df/AΔ7^*) mutants (n=3). **(C-E)** Epi-fluorescent CNS images of *babo^Df/+^*, *babo^Df/Fd4^*, and *A-indel* (*babo^Df/AΔ7^*) dissected at 120 h AEL and labeled with anti-Dlg. **(F)** A bar graph showing mean areas (± sd) of brain lobes of control (*babo^Df/+^*) and *A-indel* (*babo^Df/AΔ7^*) brain lobes at 120 h AEL, and *babo^Df/Fd4^* at 120 h and 144 h AEL. Denotations above the bars indicate statistical significances between control and mutant and otherwise indicated with brackets. Sample numbers are shown at the base of bars. Scale bars in A and C: 100 μm.

### Isoform mutants exhibit altered larval and pupal development

We investigated pan-*babo* and individual isoform mutants to further find general and/or unique roles played by the isoforms during larval and pupal development. Fresh hatchlings of *babo^Df/Fd4^* do not look noticeably different from the control (*babo^Df/+^*) in general appearance and behavior. When they were raised on a low-sugar diet to avoid early larval lethality due to sugar toxicity and competition [40], about 20% of *babo^Df/Fd4^* larvae survived until 120 h AEL (n=200). The mutant survivors move sluggishly, are about 25% shorter in body length, and exhibit conspicuously swollen anal pads (Fig 5A, 5B, 5F). Small imaginal discs and small brain phenotypes are also evident, as described previously [39]. On rare occasions, the mutant larvae develop into puparia (2 out of 200); however, they fail to evert anterior spiracles, are impaired in cuticle hardening and tanning, and die as prepupae or early pupae (Fig 5G and 5H, 5Gi and 5Hi).

**Fig 5.**
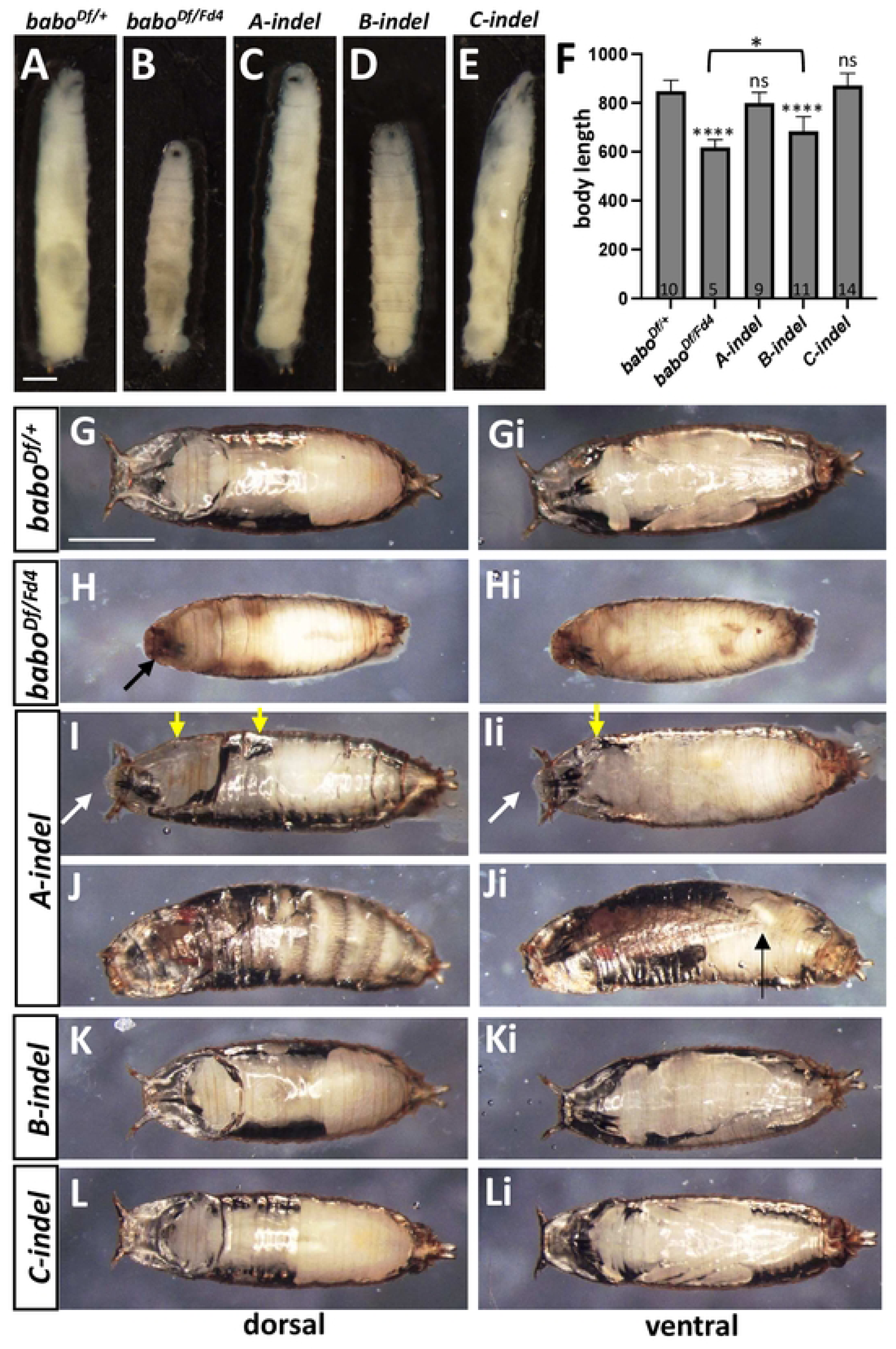
Larval and pupal development of pan-*babo* and isoform mutants at 120 h AEL. **(A-E)** Larval pictures representing *babo^Df/+^* (con)*, babo^Df/Fd4^, A-indel (babo^Df/AΔ7^*)*, B-indel (babo^Df/BΔ4^*), and *C-indel (babo^Df/CΔ^*^1^) at WL3 stage. **(F)** Mean body length (± sd) for each genotype. The number of larvae examined is shown at the base of each bar. Statistical significance between each genotype and control is indicated above the bar and, otherwise indicated with a bracket. **(G-L, Gi-Li)** Dorsal (G-L) and ventral (Gi-Li) views of pupae of indicated genotypes. (G, Gi) A control pupa. (H, Hi) A rare *babo^Df/Fd4^* developed up to the early pupal stage. A black arrow in (H) indicates failed eversion of anterior spiracles. (I, Ii) An *A-indel* mutant at early pupal stage.

Yellow arrows indicate big air bubbles under the cuticle. White arrows point to unsealed anterior part that is slightly extended. (J, Ji) The curved pharate body of an *A-indel* mutant. A black arrow indicates short legs. (K, Ki) A stout *B-indel* pupa. (L, Li) A normal-looking *C-indel* pupa. Scale bars: in A, 500 μm; in G, 1 mm.

By comparison, most *A-indel* mutant larvae develop into WL3. They show significant size reduction in both imaginal discs and brain and a slightly slender body shape but otherwise normal appearance (Fig 5C and 5F). However, they lack characteristic wandering behavior (crawling away from the food) and instead, pupariate on the food surface. Their puparium formation is abnormal; the mutant puparia fail to fully retract to a typical barrel shape, thus remaining slightly elongated and are faulty in cuticle hardening and tanning (Fig 5I and 5Ii). These phenotypes resemble those described for *myo^Δ1^* [41], supporting an intimate ligand-receptor partnership between the two. After puparium formation, many *A-indel* mutants show collapsed prepupal casings and large air bubbles trapped under the cuticle (Fig 5I and 5Ii). These features might be associated with incomplete sealing at the anterior end of the puparium which remains open until the pupal stage (Fig 5I and 5Ii). We found about 20% of *A-indel* prepupa advance to pharate adults (n=140); however, they show short legs and a curved body form and fail to eclose (Fig 5J and 5Ji).

At 120 h AEL, *B-indel* larvae are shorter than controls but still longer than pan*-babo* mutants in their body length (n=120, Fig 5D and 5F). Stout *B-indel* mutant pupae undergo normal pupal development but rarely eclose (n=120, Fig 5K and 5Ki). These phenotypes are reminiscent of those described for *Act*β-null mutants (*Act*β*^ed80^*), which is consistent with the proposed signaling of *Actβ* through BaboB [37].

Lastly, *C-indel* larvae display the most conspicuously swollen anal pads among isoform mutants but otherwise have a normal appearance in larval body length, pupal shape, and eclosion when raised on a reduced sugar diet (n=120, Fig 5E and 5F, 5L and 5Li). These phenotypes are comparable to those described for *daw* mutants, again supporting the notion that Daw signaling is primarily through the BaboC isoform [36]. The two independent *indel* mutations of each isoform gave similar results. In summary, these observations support that each isoform directs different aspects of larval and pupal development.

### Pan-*babo* and *A-indel* mutants exhibit differences in their temporal profiles of brain growth

Pan-*babo* and *A-indel* mutants differ in severity of the brain growth defect at L3 stage. To see if this trend is observed at earlier larval stages, we examined the growth of three brain regions of control (*babo^Df/+^*), pan-*babo* (*babo^Df/Fd4^*), and *A-indel* (*babo^Df/AΔ7^*) mutants at four developmental timepoints: 48 h (late L1-early L2), 72 h (Late L2-early L3), 96 h (early to mid-L3), and 120 h (WL3) AEL. While control brains increase nearly five-fold in brain lobe area from 48 h to 120 h AEL (Fig 6A-6D), pan*-babo* mutants show retarded growth in both OL and CB even at 48 h AEL (Fig 6A). In contrast, brain regions of *A-indel* mutants are not significantly different at 48h AEL, suggesting relatively normal development in the absence of BaboA function during early stages (Fig 6A). The *A-indel* mutants show obviously retarded growth in the OL at 72 h AEL (Fig 6B) but not in the CB until 96 h AEL (Fig 6C). Taking these results together suggests that pan-Babo function is required for proper growth of both the OL and CB regions from early to late larval stages, while BaboA function becomes critical for proper growth of the OL starting from sometime between 48 to 72 h AEL and the CB, between 72 and 96 h AEL. We also point out that both mutants do not show significant size difference in the CB at 120 h AEL, suggesting that differential OL development primarily accounts for the brain size disparities between pan-*babo* and *A-indel* mutants at late larval stage.

**Fig 6.**
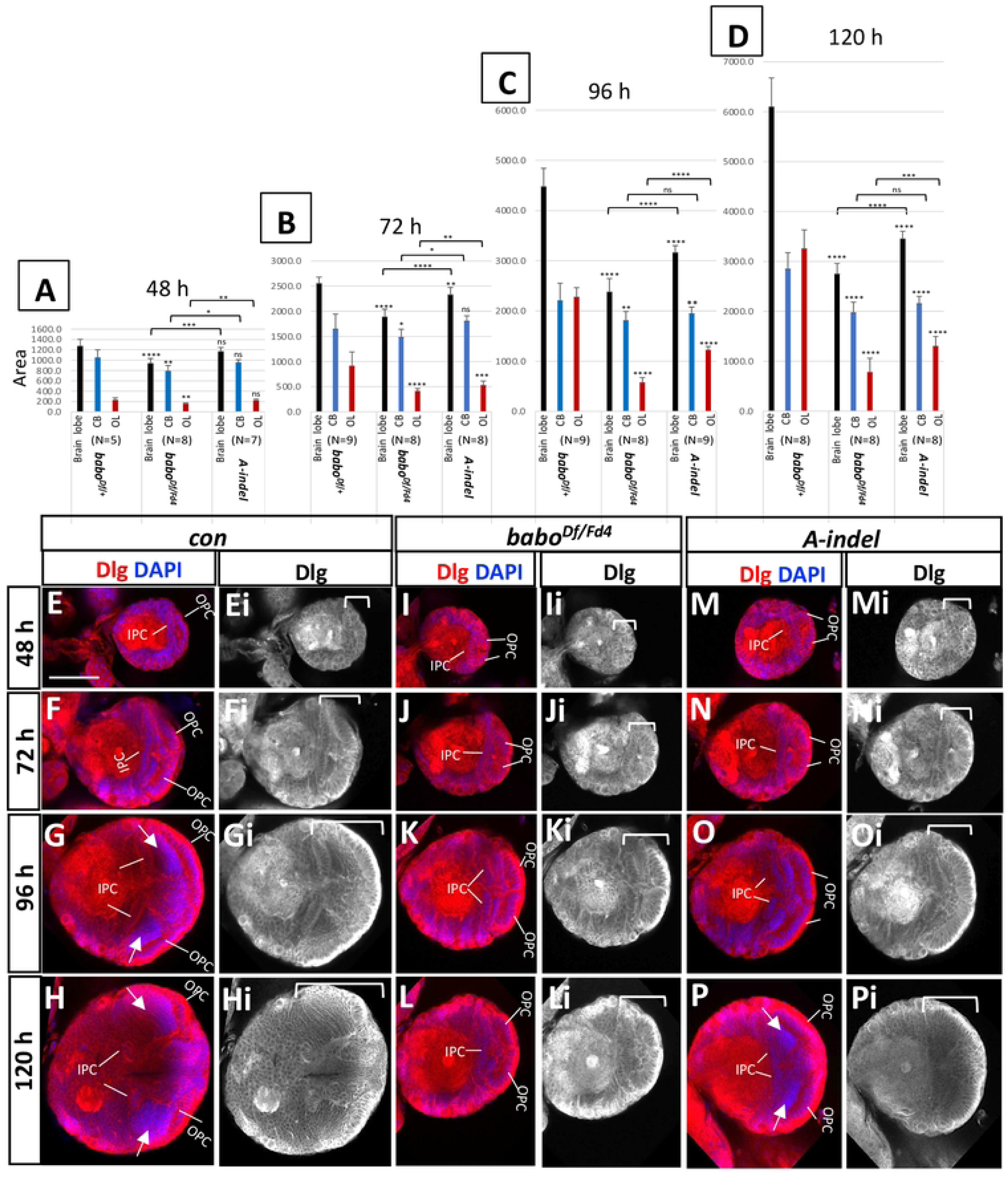
Growth defects in the CB and OL regions of pan-*babo* and *A-indel* mutants. **(A-D)** Mean areas (± sd) of three regions, whole brain lobe (black), CB (blue), and OL (red) of control (*babo^Df/+^*), pan-*babo* (*babo^Df/Fd4^*) and *A-indel* (*babo^Df/AΔ7^*) mutants at 48h (A), 72 h (B), 96 h (C), and 120 h (D) AEL. Statistical significance between each genotype and con is indicated above the bars or otherwise indicated with brackets. **(E-P, Ei-Pi)** Frontal mid-sections of brain lobes showing merge of Dlg (red) and DAPI (blue) staining (E-P), or Dlg staining alone (grey, Ei-Pi). Medulla field is indicated by white arrows (G,H,P). Brackets show the span of the OL. Scale bar in E: 50 μm.

### Defective OL development of pan-*babo* and *A-indel* mutants

Since our foregoing results show that reduced OL growth largely accounts for brain size difference between pan-*babo* and *A-indel* mutants, we monitored the development of this primordial tissue more closely. The NE cells in the OPC and IPC can be distinguished by their characteristic columnar cell shape, arrangement, and position. The control OL at 48 h AEL is composed exclusively of NE cells that are organized into two well-defined layers each representing OPC and IPC (Fig 6E and 6Ei). Both proliferation centers greatly increase their population by symmetric divisions of NE cells (Fig 6F and 6Fi). At 96 h AEL, a densely DAPI-stained area found in the medulla field between the OPC and IPC denotes a population of newly generated medulla precursor neurons, indicating ongoing NE-to-NB conversion as well as rapid proliferation of the resulting NBs in the OPC (Fig 6G and 6Gi). At 120 h AEL, NE populations in both proliferation centers have declined considerably due to continuous transformation of NEs (Fig 6H and 6Hi). These observations are consistent that an increasing NE population accounts for the growth of this region until 72 h AEL and afterward the growth is fueled by a growing population of precursor cells generated by rapidly proliferating NBs.

When compared to control, pan-*babo* mutants have fewer NE cells at 48 h AEL (Fig 6I and 6Ii). The NE population of the mutants appears to increase at a slower pace up to 96 h AEL (Fig 6Ii-6Ki) and are mostly devoid of DAPI-stained precursor cells in the presumptive field even at 120 h AEL (Fig 6L and 6Li). Thus, the overall appearance and size of the pan-*babo* OLs of 120 h AEL are not so different from theirs at 96 h AEL, which show close resemblance to the control OLs of 72 h AEL, suggesting that slower proliferation of NEs and poor generation of medulla precursor neurons, together contributing to the severely reduced OL growth in mutant. The latter event might explain the lack of growth in the mutant OLs from 96 h to 120 h AEL (Fig 6C and 6D, 6K and 6L). These observations support the notion that pan-Babo function is required for regulating both proliferation of NEs and production of medulla precursor neurons. Unlike the pan-*babo* mutants, *A-indel* OL primordia display normal-looking NEs in both size and number of at 48 h AEL (Fig 6M and 6Mi) but a noticeably smaller NE population at 72 h AEL (Fig 6N and 6Ni), consistent with the timing of significant reduction first detected in this area (Fig 6B). These results together with substantially fewer DAPI-stained medulla precursor neurons observed at 96 h and 120 h AEL (Fig 6O and 6P, 6Oi and 6Pi) indicate that late proliferation of NEs and production of medulla precursor neurons are also compromised in *A-indel* mutants.

### Slow proliferation of two NB types in the CB of pan-*babo* and *A-indel* mutants

The larval CB contains two NB types, type-I and type-II. More common type-I NBs divide asymmetrically to produce a series of ganglionic mother cells (GMCs), each of which divides once to give rise to two precursor cells. The type-II NBs also divide asymmetrically but generate intermediate neural progenitor cells (INPs) that act like temporary NBs producing several GMCs [42]. Both pan-*babo* and *A-indel* mutations show significant size reduction in the CB region, possibly due to attenuated mitotic activities of NBs. To address this, we examined NBs and their lineage cells labeled with anti-Dlg. While Dlg expression is detected in all NBs and their clones, staining intensities are noticeably stronger in NBs, GMCs, and recently born precursor cells, which allows us to gauge mitotic activities of NBs. We focused on a group of dorsal NBs in the CB. Control NBs display a large population of NB-derived precursor cells with intense Dlg staining (Fig 7A), indicative of their robust ongoing proliferation. In contrast, pan-*babo* mutant NBs are associated with far fewer precursor cells displaying intense Dlg immunostaining (Fig 7B), while *A-indel* mutants show intermediate numbers of such cells between control and pan-*babo* mutants (Fig 7C).

**Fig 7.**
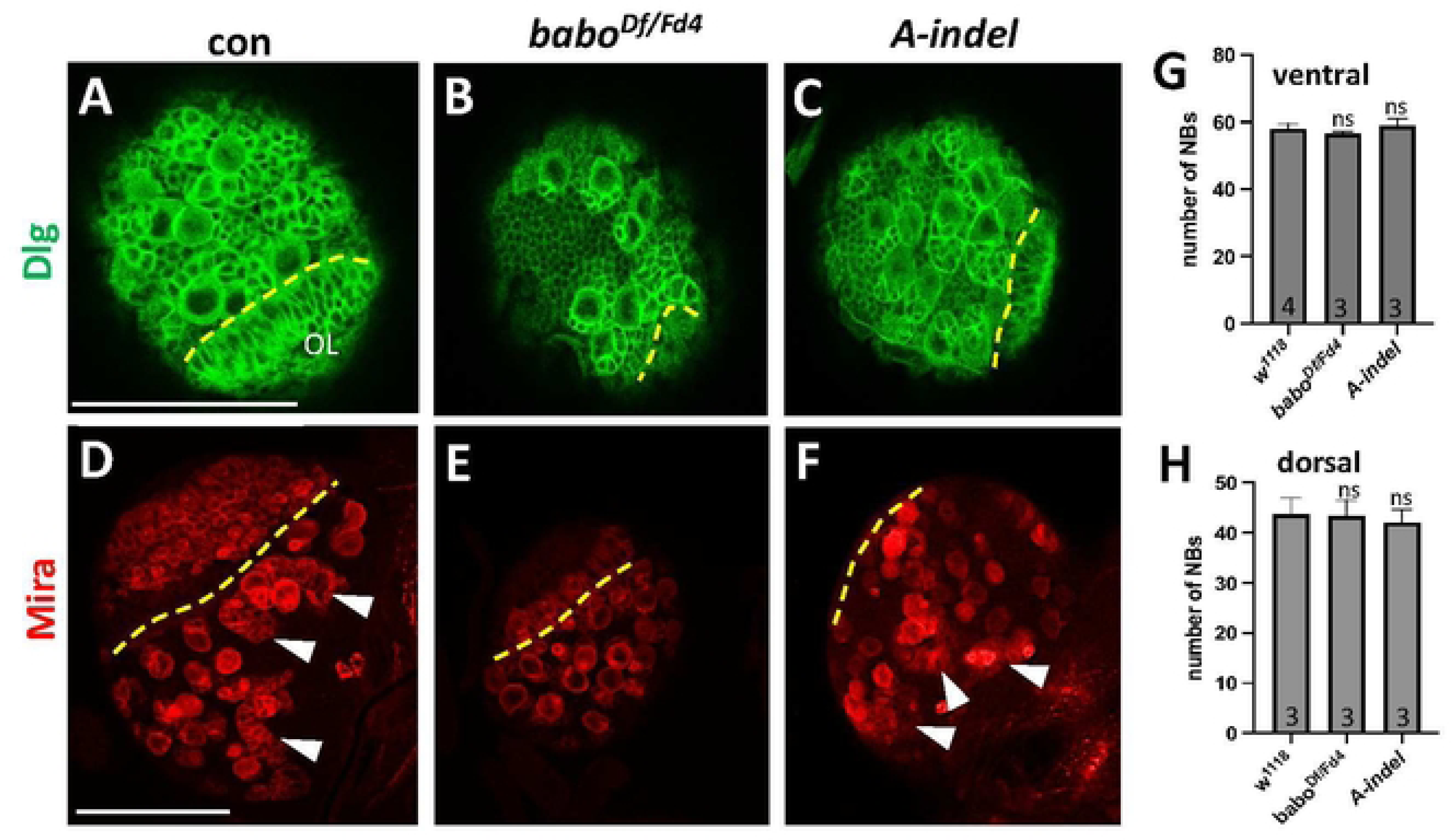
Retarded proliferation of NBs in the CB of pan-*babo* and *A-indel* mutants at 120 h AEL. Dotted lines mark the boundary between the OL and CB. **(A-C)** Confocal images showing the dorsal NBs and their clones labeled with anti-Dlg (green) of *babo^Df/+^* (con), *babo^Df/Fd4^*, and *A-inde* (*babo^Df/AΔ7^*). NBs and late born precursor cells are stained strongly. **(D-F)** Anti-Mira (red) labels NBs and most recently born precursor cells. Three type-II NBs and their tailing clones are indicated by arrowheads. **(G,H)** Mean numbers of NBs (± sd) counted in ventral and dorsal sides of brain lobe. Scale bars in A and D: 50 μm.

We also examined the mitotic activity of dorsal NBs using anti-Mira that labels NBs, INPs, GMCs, and only late-born precursor cells. Most of the control type-I NBs abut at least one GMC and several small progeny cells in the corner of their somata (Fig 7D and S2A). In contrast, pan-*babo* mutant display type-I NBs with significantly fewer Mira-positive clones (Fig 7E and S2B). *A-indel* mutants show intermediate numbers of Mira-positive clones accompanying type-I NBs when compared to control and pan-*babo* ones (Fig 7F and S2C), confirming sluggish proliferation of type-I NBs in pan-*babo* and *A-indel* mutants.

Additionally, the mitotic activities of type-II NBs were assessed in these two. Each *Drosophila* larval brain lobe contains eight type-II NBs in the dorsal and six in the medial region. In the control, a string of Mira-positive cells trails individual type-II NBs, a sign of ongoing fast divisions (Fig 7D) but such characteristic trails are not observed in pan-*babo* mutants, indicating idling proliferation in mutant type-II NBs (Fig 7E). *A-indel* mutants show slow proliferation of type-II NBs too but not to the extent of pan*-babo* mutants (Fig 7F). Based on these results, we conclude that pan-Babo function is needed to either promote or maintain a proper mitotic activity of both NB types and BaboA function is primarily but not exclusively responsible for this activity, thus expecting involvement of other isoforms. We also counted the number of Mira-positive NBs on the ventral and dorsal sides of the CB at the WL3 stage to determine if the reduction in the CB area also results from fewer NBs. This appears not to be the case, since the total numbers of NBs from both pan-*babo* and *A-indel* mutants are the same as control ones (Fig 7G and 7H, S2). Henceforth, we argue that the smaller CB of *babo* mutants results not from a smaller NB population, but from their reduced proliferation.

### BaboA acts in a canonical TGF-β signaling pathway through dSmad2

Previous studies indicate an intimate Myo-BaboA partnership for regulating imaginal disc size [38], remodeling of the MB γ neurons [41], and apoptosis of the ventral corazonergic neurons [43]. In line with these observations, we found that A*-indel* and *myo^τ1^* mutants phenocopy each other in many other aspects of larval and pupal development (Fig 5). Moreover, the two genotypes exhibit very similar growth defects in the CB and OL at 120 h AEL, again supporting the Myo-BaboA relationship for postembryonic brain growth (Fig 8A).

**Fig 8.**
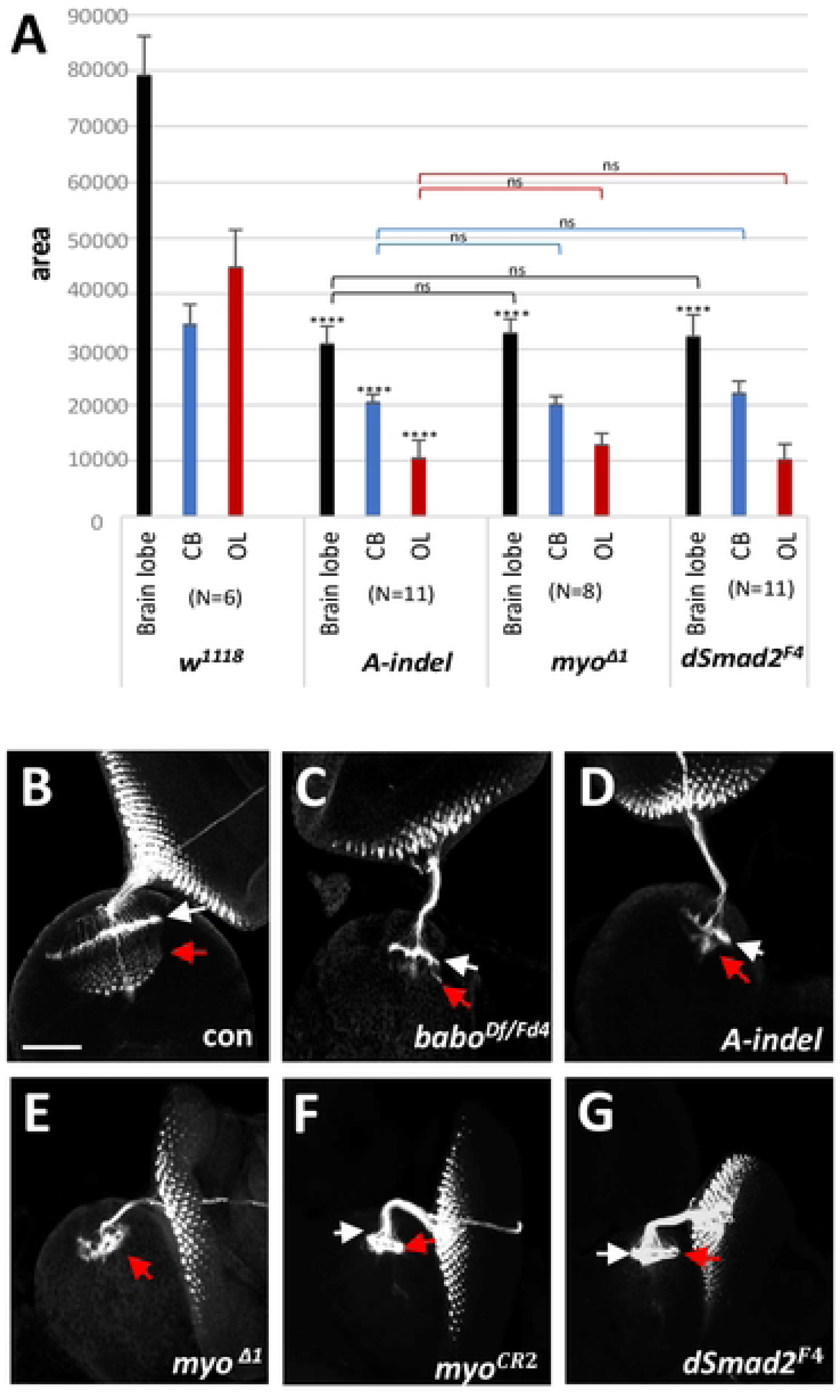
BaboA acts through the canonical TGF-β signaling pathway for growth of brain lobe regions. **(A)** Mean areas (± sd) of three brain lobe regions of *w^1118^*(con), and mutants homozygous for *A-indel* (*bab ^AΔ7^*), *myo^Δ1^*, and *dSmad2^F4^* (*dSmad2^F4^*/Y) at 120 AEL. Statistical significance is indicated above each bar or bracket. (ns, not significant; N, sample numbers). **(B-G)** Retinal photoreceptor neurons and their axon bundles labeled with anti-24B10 in the developing OL of *babo^Df/+^* (con, n=14), *babo^Df/Fd4^* (n=6), *A-indel* (*babo^Df/AD7^*, n=12), *myo^Δ1^* (*myo^Δ1/Δ1^*, n=6), *myo^CR2^*(*myo^CR2/CR2^*, n=5), and *dSmad2^F4^* (*dSmad2^F4^*/Y, n=5). All animals were dissected at 120 h AEL, except for those of *babo^Df/Fd4^*that were dissected at 144 h AEL. White arrows indicate the lamina plexuses, and red arrows, axonal innervations in the medulla area. Scale bars in A: 50 μm.

The canonical downstream effector of Babo is dSmad2 (Smox), the only R-Smad of Activin-type TGF-β signaling in *Drosophila*. However, it has been shown that Babo can also signal non-canonically to stimulate growth of the wing disc [39]. To see which is the case for brain growth, three brain areas were compared between *A-indel* and *dSmad2^F4^*, a null allele that deletes the entire *dSmad2* coding region [44]. Since dSmad2 is a common downstream effector in the canonical TGF-β signaling for all three Babo isoforms, we expected that the small brain phenotype of *dSmad2^F4^* would resemble that of pan-*babo* mutants. To our surprise, mean areas of the brain lobe, CB, and OL of *dSmad2^F4^* do not noticeably differ from those of *A-indel* and *myo^ι1^*mutants (Fig 8A). It also means that they are larger than the sizes of the same structures in pan-*babo* mutants. These results suggest that BaboA works through the canonical TGF-β signaling pathway to promote the growth of the CB and OL, whereas the more severe size reduction found in the pan*-babo* mutant suggests that brain growth regulated by other Babo isoforms involves a dSmad2-independent pathway.

### pan-*babo* and *A-indel* mutations similarly disrupt retinal axon targeting

The *Drosophila* compound eye consists of 750-800 ommatidia, each containing eight photoreceptor neurons (R1-R8). The axons of R1-R6 terminate in the lamina forming an ostensive neural plexus, whereas the axons of R7 and R8 extend into the medulla field establishing intricate patterns of synapses with developing medulla neurons [45–47]. Given under-development of the pan-*babo* mutant optic field, we examined the retinal axon targeting patterns using anti-24B10 (Chaoptin). In pan*-babo* mutants, the retinal axon bundles track normally through the optic stalk but subsequent targeting in both the lamina and their medulla fields is severely impaired (Fig 8B vs. 8C), consistent with previous reports with other *babo* alleles [34,35]. It is not better in *A-indel* mutants (Fig 8C vs. 8D). This is surprising since they contain substantially more medulla precursor neurons in their presumptive medulla field compared to pan-*babo* mutants (Fig 6L vs. 6P). Similarly defective retinal axon targeting is also found in two different *myo*-null mutants, *myo^τ1^* and *myo^CR2^*(Fig 8E and 8F) as well as in *dSmad2^F4^* mutants (Fig 8G). These results not only reinforce the idea that Myo-BaboA-dSmad2 signaling is required for the proper generation of precursor neurons but also suggest its key role in differentiation of lamina and medulla neurons within the OL.

### Defective NE conversion in pan-*babo* and *A-indel* mutants

Prior to mid-L3, NEs in the OPC begin their transformation into medulla NBs medially and LPCs laterally, which then generate medulla and lamina precursor neurons, respectively [24,48]. To see if the poorly developed medulla fields of pan-*babo* and *A-indel* mutants are due to compromised NE conversion, we examined medulla NBs with anti-Mira and LPCs and lamina precursor neurons with anti-Dachshund (Dac) at WL3 stage. The medulla NBs are readily distinguishable from ventrally located CB-NBs based on their size and location. In the *w^1118^* control, we observe a large population of Mira-positive medulla NBs present in multiple layers, as well as numerous Dac-positive LPCs and lamina precursor neurons (Fig 9A and 9Ai). In stark contrast, both pan-*babo* and *A-indel* mutants contain much fewer layers of medulla NBs (Fig 9B and 9C), although *A-indel* mutants have slightly more than pan-*babo* mutants, which is consistent with *A-indel* mutants having an intermediate phenotype. Similar trend is seen in Dac-positive LPCs (Fig 9Ai-9Ci). These results explain the smaller medulla precursor populations observed in pan-*babo* and *A-indel* mutants, respectively (Fig 6L and 6P). The combined observations suggest that not only the proliferation of NEs but also the conversion of NEs to medulla NBs and LPCs are defective in both mutants.

**Fig 9.**
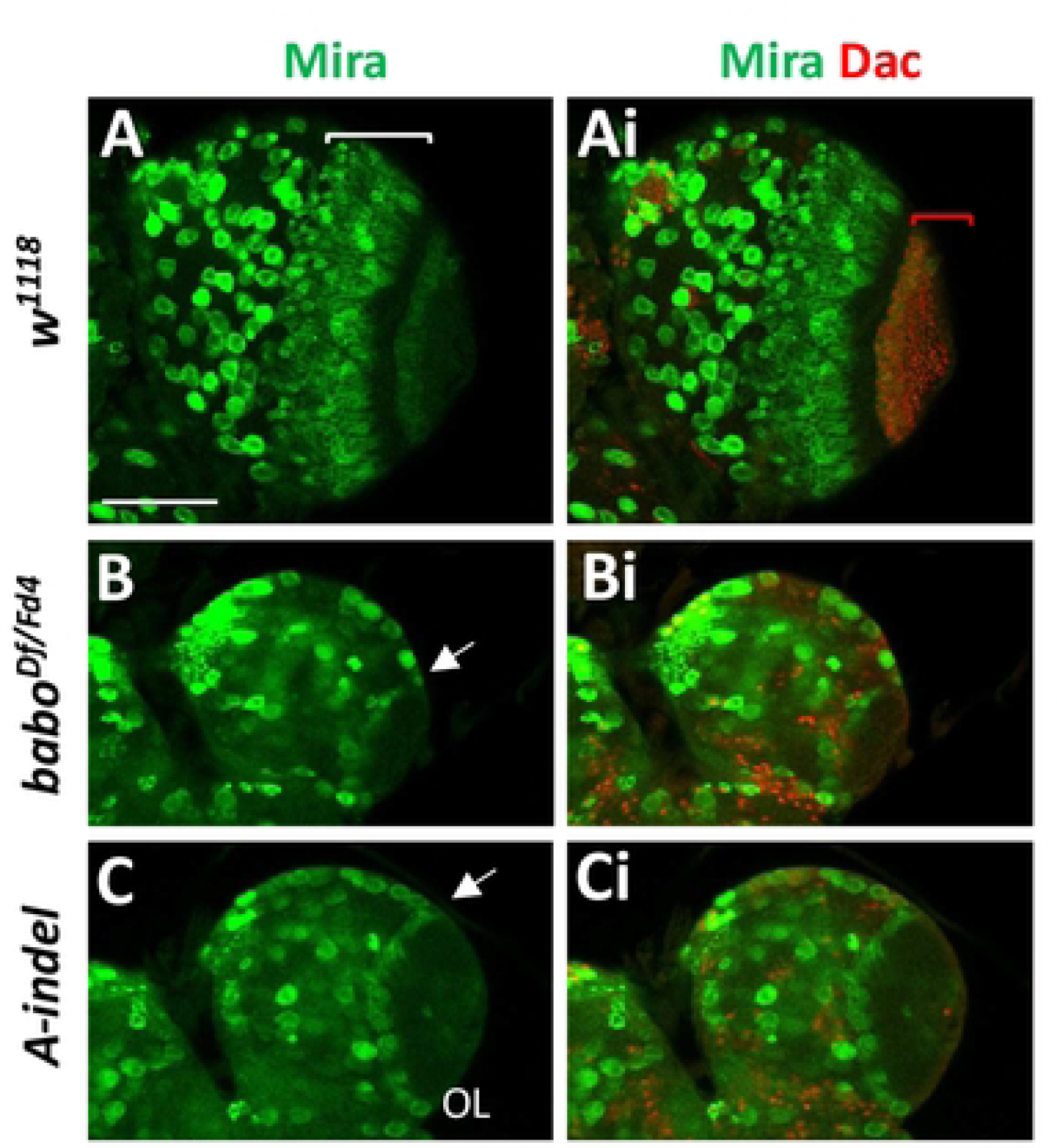
Severe reduction of medulla NBs and LPCs in pan-*babo* and *A-indel* mutants. Ventral OL views at WL3 stage showing medulla NBs labeled with anti-Mira (green) and lamina precursor neuronal cells (LPCs) labeled with anti-Dac (red). **(A,Ai)** *w^1118^* (con, n=7). A span of layers of medulla NBs is indicated by a white bracket and Dac-positive cells by a red bracket. **(B,Bi)** *babo^Df/Fd4^* (n=6). **(C,Ci)** *A-inde*l (*babo^Df/AD7^*, n=7). Mira-positive medulla NBs in a layer are indicated by arrows. Scale bar in A: 50 μm.

### Autonomous requirement of *baboA* in NEs

To confirm the autonomous action of BaboA in NE and NB cells, we tested whether knockdown (KD) of *baboA* expression specific to the two cell types phenocopies *A-indel* mutants. We employed the *R31H09-gal4* that drives reporter expression strongly in the NEs of both OPC and IPC and relatively weakly in CB-NBs during larval development [49]. *R31H09-gal4* flies were crossed to *UAS-baboA-*miRNA (*baboA*-KD) or *w^1118^* (con) flies and brain growth was assessed in their progeny at the WL3 stage. *baboA*-KD causes significant brain hypotrophy, which is primarily the result of a ∼50% reduction in the OL (Fig 10A). No significant size changes are seen in the CB, presumably due to insufficient knockdown efficacy in CB-NBs. Nevertheless, these data strongly support an autonomous role for BaboA in proliferation and/or conversion of NEs.

**Fig 10.**
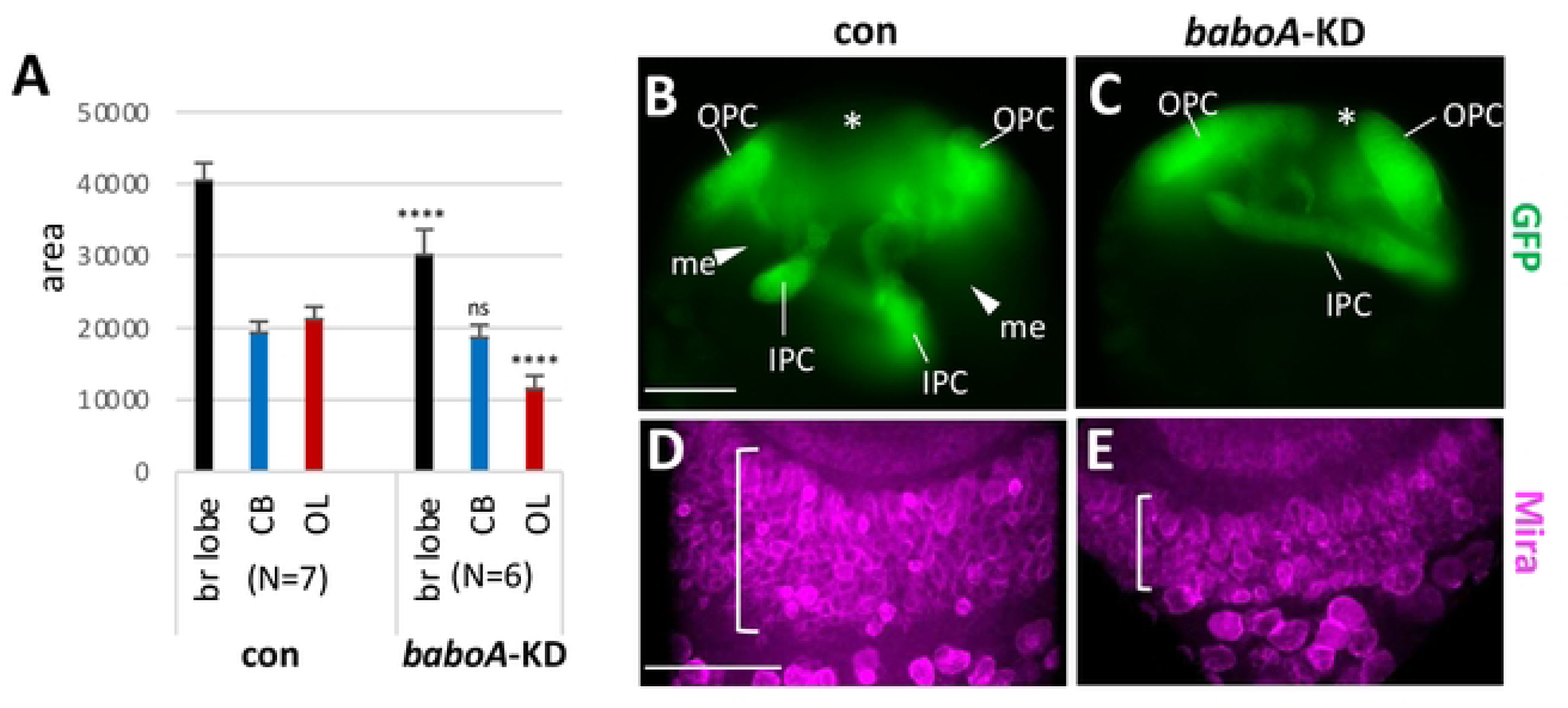
*baboA* knockdown (KD) in NEs causing reduction in medulla NB population and optic lobe area. *UAS-mCD8GFP;R31H09-gal4* flies were crossed to *w^1118^* (con) or *UAS-baboA-miRNA* for *baboA*-KD, respectively. Progeny brains were examined at WL3 stage. **(A)** Mean areas (± sd) of the three brain regions. **(B,C)** mCD8GFP (GFP) expression is detected strongly in NEs of both genotypes. The medulla field (me) is indicated by arrowheads and the center of distal-IPC zone by asterisks. **(D,E)** *baboA*-KD in OPC NEs reduces significantly the numbers of Mira-positive (magenta) medulla NBs. A population of medulla NBs on the ventral brain region are indicated by brackets. Scale bars in A and D: 50 μm.

Next, we examined GFP expression in the frontal mid-sections of the OL in control and *baboA*-KD at the WL3 stage. Intense GFP signals mark small strips of OPC and IPC NEs (Fig 10B). A decreased population of OPC NEs and a wide space between IPC and OPC layers denote ongoing NE-to-NB conversion followed by active proliferation of medulla NBs. In contrast, *baboA-*KD brains show long strips of GFP-positive NE cells from both proliferation centers that are still is close apposition (Fig 10C); such a cellular organization resembles that of *A-indel* mutants (Fig 6Oi and 6Pi). In addition, a large population of medulla NBs labeled with anti-Mira is shown on the ventral side of control OLs, indicative of actively ongoing NE-to-NB transformation, while *baboA-*KD exhibit a smaller medulla NB population (Fig 10D and 10E). A large gap is also noted between OPC strips, which is the result of both dwindling OPC NE population and rapid growth of both the lamina field and distal-IPC (d-IPC) zone (Fig 10B). The latter consists of IPC-originated NBs and their clones, which later give rise to the lobula plate neurons [50]. In contrast, the gap is not prominent in *baboA*-KD, indicating that both lamina and the d-IPC zones are not properly developing (Fig 10C). These results, together with intensive GFP expression in IPC NEs, led us to propose that transformation of IPC NEs is also compromised by *baboA-KD*. However, since *R31H09-gal4* is active in NEs from early larval stages. it is not entirely clear if the NE conversion event is preordained by BaboA functioning in earlier NEs or needs it for this event too.

### BaboA function is needed for proper NE conversion

To address when BaboA expression is required in the NE for proper NB conversion, we crossed *tub-gal80^ts^; da-gal4* flies to *w^1118^* (control), *UAS-baboA-miRNA* (*baboA*-KD), or *UAS-dSmad2-RNAi* (*dSmad2*-KD) flies. The resulting progeny were raised at 18°C until they started to show the wandering behavior and afterward at 29°C until adult emergence. Unlike *A-indel* mutants, the *baboA*-KD larvae develop into normal-looking adults, implying no detrimental effect on pupal development and eclosion (Fig 11A vs. 11B, 11D vs. 11E). However, eclosed *baboA*-KD adults are shorter and with slightly upraised wings (Fig 11E and 11I). The *dSmad2*-KD also show normal-looking pupal development but fail to eclose (Fig 11C). Thus, we used the pharate brains of the three genotypes to measure regional areas. Both *baboA*-KD and *dSmad2*-KD brains similarly show a nearly 50% reduction in the OL region but no significant change in the CB area (Fig 11F-11H, 11J). These results are consistent with the OL region continuously increasing its size during and after the WL3 stage [32,51]. We also reasoned that CB growth is less affected by the knockdown of *baboA* or *dSmad2* from this period on, with the caveat that the knockdown may not be complete in CB-NBs.

**Fig 11.**
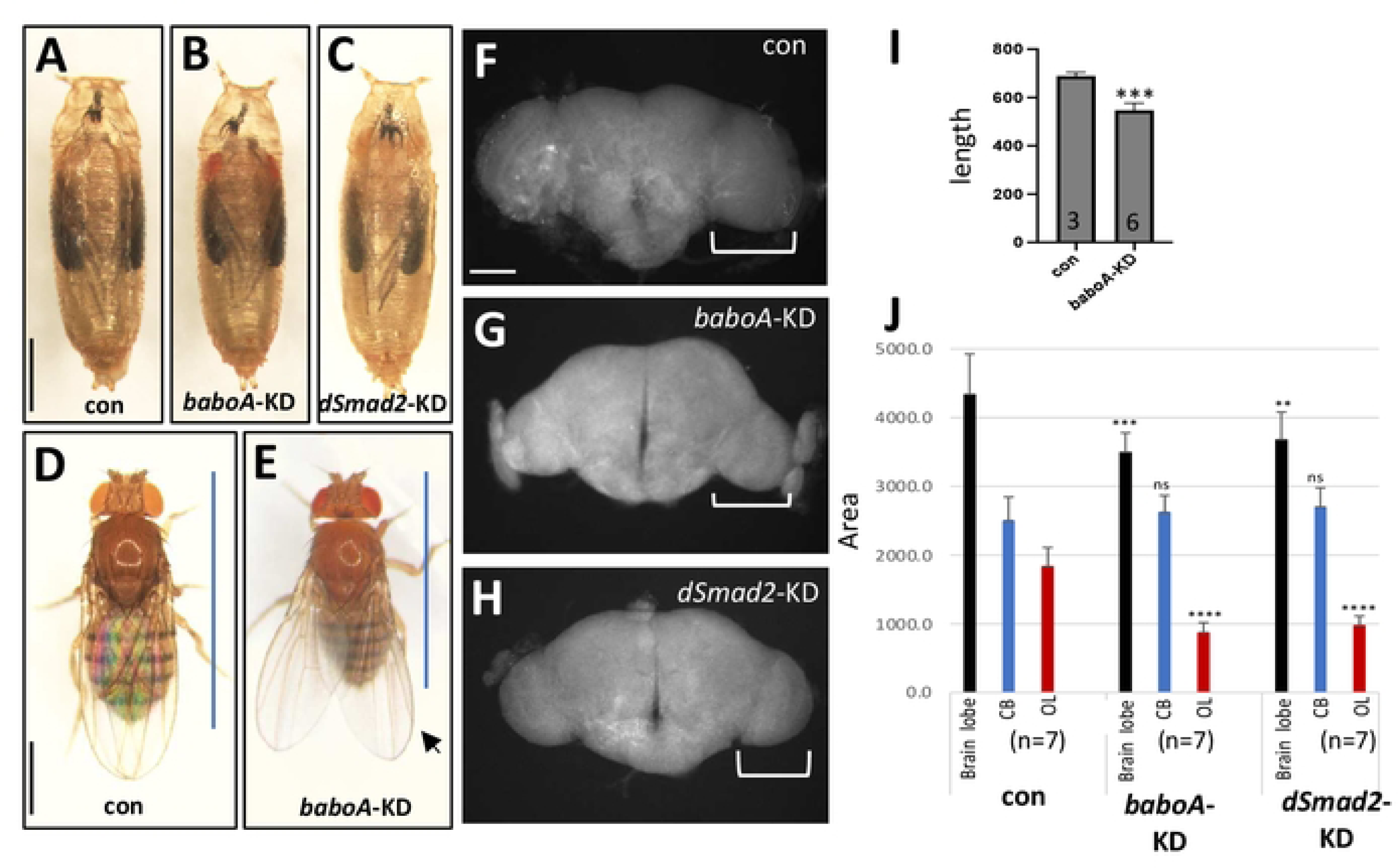
Stage-specific knockdown effect of *baboA* and *dSmad2* on adult body and brain size. *tub-gal80^ts^; da-gal4* flies were crossed to *w^1118^*, *UAS-baboA-miRNA*, and *UAS-dSmad2-RNAi* for con, *baboA-KD*, and *dSmad2-KD*, respectively. Laid eggs (F1 progeny) were raised at 18°C up to early WL3 stage and then at 29°C until adult emergence. **(A-C)** Pharate bodies of con (n=40), *baboA*-KD (n=30), and *dSmad2*-KD (n=30). **(D,E)** Adults of con and *baboA*-KD. Blue lines indicate span of body length, and an arrow points to slightly upraised wings. **(F-H)** Epi-fluorescent brain images of con (n=7), *baboA*-KD (n=8), and *dSmad2*-KD (n=7) pharate adults. The optic lobe is indicated by brackets. **(I)** A graph showing shorter body length of *baboA*-KD adults. Only females were used for this analysis. **(J)** Mean areas (± sd) of the brain lobe (black), CB (blue), and OL (red) of pharate brains. Statistical significance with con is indicated above bars. Scale bars in A and D, 1 mm and in F, 100 μm.

Since the NEs stop their proliferation around the mid-L3 stage, two of the key cellular events that can affect the growth of the OL are NE conversion and proliferation by OL-NBs, both of which are observed until the end of the prepupal stage, around 12 h APF [51]. To test if BaboA function is also needed for OL-NB proliferation, *baboA*-KD was induced by a *scro*-*gal4* that is active only in a subpopulation of medulla NBs, and continuously in their clones, but not in OPC NEs [5,52]. *UAS-mCD8GFP; scro-gal4* was crossed to *w^1118^* (control) or *UAS-baboA-miRNA* and GFP expression was examined on the ventral side of progeny brains at the WL3 stage. We found that *baboA*-KD does not considerably reduce the combined population of both *scro*-positive cell types within the OPC zone (S3 Fig). These results imply that *baboA*-KD in medulla NBs does not greatly affect the proliferation event of medulla NBs, further suggesting that the primary cellular event compromised by stage-specific *baboA*-KD is NE-conversion event. Therefore, we conclude that BaboA function is needed separately for the two cellular events of NE cells: promoting conversion and proliferation.

### Babo isoforms are epistatic to neural expression levels of EcR-B1

Previous studies have shown that EcR-B1 expression is regulated in the NEs and NBs in a stage-specific manner [21,32] and disruption of its function by overexpressing a dominant negative form of EcR (EcR-DN) blocks transformation of OPC NE-to-NB and CB-NB proliferation rate [21, 52]. As expected, *R31H08>EcR-DN* causes noticeably smaller brains due to significant reduction in both OL and CB (S4 Fig). It is notable, however, that this is more severe effect than *R31H08>baboA-miRNA* which does not cause substantial decrease in the CB region. We also found that EcR-DN expression severely blocks NE-to-NB conversion in the OPC and IPC, which is again more severe than that of *baboA-KD* (S4 Fig). In general, the *R31H08>EcR-DN* closely phenocopies the pan-*babo* null mutation (S4 Fig).

High levels of EcR-B1 are detected in various cell types in the CNS including NEs and NBs at the WL3 stage [21, 32, 53]. Such EcR-B1 expression is barely detectable in the CNS of pan-*babo* mutants, confirming an essential function of Babo for the stage-specific upregulation of EcR-B1 expression in this organ [18]. The canonical TGF-β signaling mediated by Myo/BaboA/dSmad2 has been shown to enhance EcR-B1 expression in the MB neurons at WL3 [18, 41]. To see if BaboA functions promote EcR-B1 expression in other cell types besides MB neurons, we examined anti-EcR-B1 immunosignals in the CNS of control (*w^1118^*) and *A-indel* mutants at 120 h AEL and pan-*babo* mutants at 144 h AEL to compensate for retarded larval development. While high levels of EcR-B1 are detected in numerous cell types including MB neurons in the control CNS (Fig 12A and 12Ai, 12D and 12Di), the levels are very low in the pan-*babo* mutants throughout the CNS except for glial cell types on the surface that show slightly more expression (Fig 12B and 12Bi, 12E and 12Ei). EcR-B1 expression levels in *A-indel* mutants are substantially lower than those in controls but still considerably higher than those in pan-*babo* mutants (Fig 12C and 12Ci, 12F and 12Fi). Our studies consistently show that *A-indel* mutant phenotypes are less severe than those of pan-*babo* mutants in all aspects of brain development and EcR-B1 levels also follow this trend. In summary, we propose that the BaboA plays a foremost role in upregulating stage-specific EcR-B1 expression in neural stem cells but that the action of additional isoforms is necessary to achieve a full degree of EcR-B1 expression either acting autonomously or not, which in turn promotes behavior changes of neural stem cells in response to increased ecdysone levels at least during the last larval stage.

**Fig 12.**
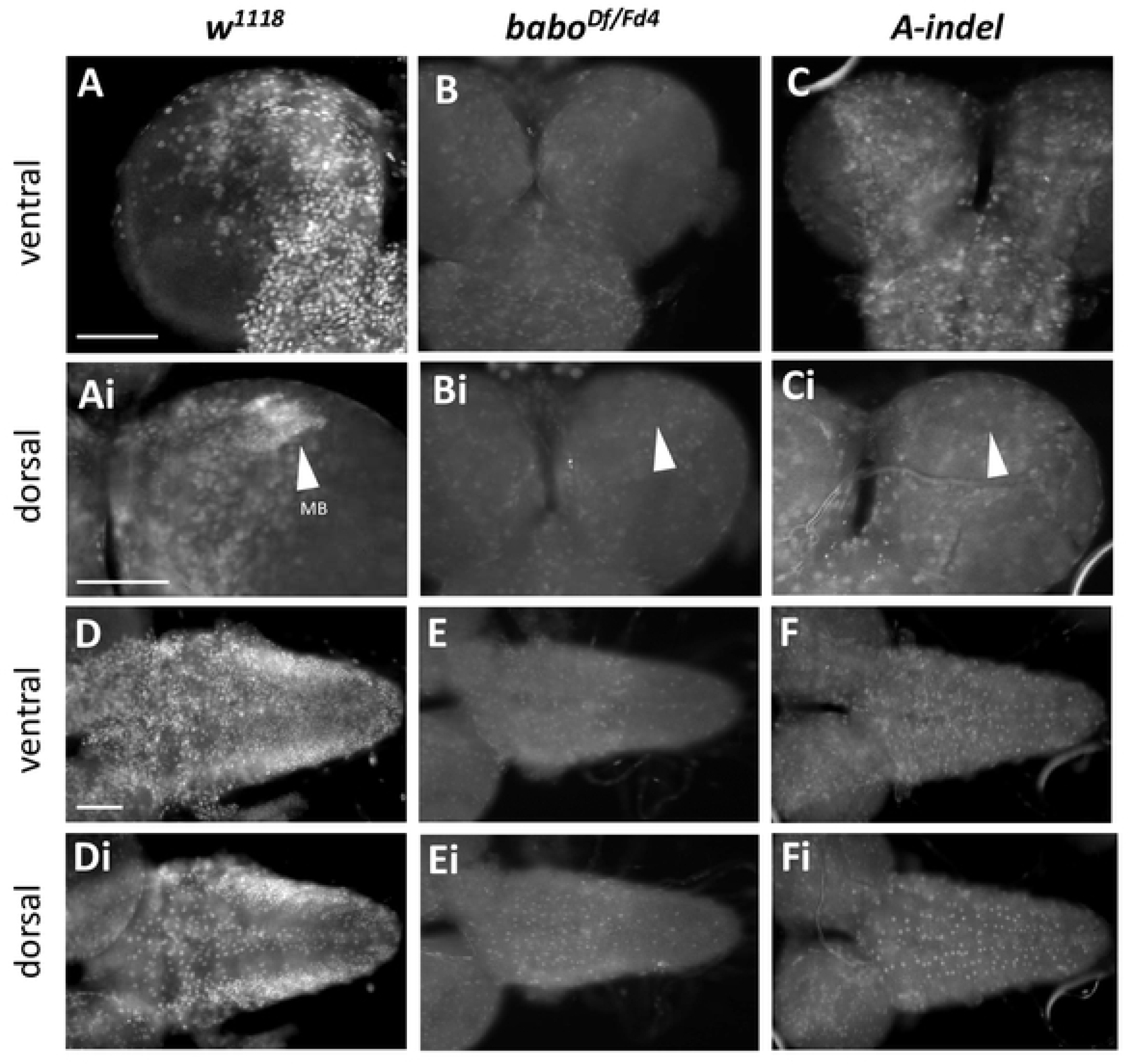
Reduced EcR-B1 expression levels in the CNS of *babo^Df/Fd4^* and *A-indel* mutants. Larval CNSs of *w^1118^* (n=14) and *A-indel* (*babo ^AΔ7/AΔ7^*, n=14) were dissected at 120 h AEL and *babo^Df/Fd4^*(n=6) at 144 h AEL to compensate for retarded development. The samples were simultaneously processed with anti-EcR-B1 and imaged under the same condition. **(A-C)** Ventral brain lobe views of indicated genotypes. **(Ai-Ci)** Dorsal view of the same samples shown in (A-C). Densely clustered mushroom body (MB) neurons are indicated by arrowheads. **(D-F, Di-Fi)** The VNCs of indicated genotypes are shown in ventral (D-F) and dorsal (Di-Fi) sides. Scale bars in A, Ai, and D: 50 μm.

## Discussion

In this study, we uncovered crucial roles of the TGF-β type-I receptor Babo in two major cellular events of postembryonic neurogenesis, proliferation of CB-NBs and OL-NEs and NE conversion, together largely contributing to dramatic larval brain growth. These events are regulated primarily, but not exclusively, by the BaboA isoform as pan-*babo* mutants exhibit more severe brain growth defects than *A-indel* mutants, indicating additional contribution of the other isoforms. In line with this, previous study with *daw*;*actβ* double mutants showed that BaboB and BaboC redundantly act in regulating brain growth [35]. However, it remains uncertain whether these two are required in the brain or in other tissues such as the prothoracic gland [54]. In the canonical TGF-β signaling dSmad2 acts as the converging downstream effector. Nevertheless, *dSmad2-*null mutants show a small brain phenotype comparable to *A-indel* and *myo*-null mutant ones which is decidedly not as strong as the pan *babo* mutant, suggesting that Myo-BaboA act through the canonical signaling pathway and the other isoforms, in a non-canonical manner. A non-canonical role of *babo* has been suggested previously in wing imaginal disc size control [39]. In this study, the removal of dSmad2 function leads to ectopic phosphorylation of Mad, the normal BMP responsive R-Smad. If a similar mechanism operates in the brain, then loss of dSmad2, but not Babo, might produce an ectopic pMad signal that leads to a low level of compensatory proliferation. In summary our studies clearly indicate that the dramatic size increase of *Drosophila* brain observed during larval stages is mediated through multiple Babo isoforms that control the behaviors of two key postembryonic neural stem cells in coordination with developmental status.

### Functions of Babo isoforms needed for proper proliferation of CB NBs

Postembryonic NBs in the central brain differ in their proliferation behavior before and after the CW checkpoint [7]. Prior to the CW checkpoint (i.e., first phase), NBs divide slowly and their division rate is sensitive to dietary protein contents, while NBs post CW checkpoint (second phase) divide faster in a manner independent of dietary status [16]. Our observations suggest that the activities of multiple isoforms are required for promoting proper proliferation of NBs in both phases, and that BaboA is acting primarily in the second phase. Since the NB proliferation in the first phase is dependent on a protein-rich diet [11,33], Babo might function in relaying feeding status to NBs. This notion is reasonable when we consider that Daw acting through BaboC is a major regulator of systemic metabolic homeostasis and cellular metabolism [40] and *C-indel* mutants display delayed larval development. Intriguingly, the timing of BaboA action in the second phase is roughly coincident with *myo* expression in the CNS [41]. One of the major *myo-*expressing cells in the central brain is the cortex glia [41], which form a specialized niche for individual NB lineages and influence NB entry into the second phase or maintenance of rapid proliferation during the second phase [4,55]. Such intercellular communication nicely aligns with the timing of BaboA action. In this manner, Babo isoforms may play important roles in providing NBs with proper developmental cues to coordinate their proliferation with their developmental status.

### Roles of pan-Babo and BaboA in the optic lobe

The behavior of OL-NEs also changes during development: expansion of the NE population by symmetric division in the first phase followed by their conversion into NBs and rapid divisions of NBs in the second phase [10,33,50]. Like NBs in the CB, the phase transition of the NEs is known to be associated with the CW checkpoint [11,33]. Pan-*babo* mutants already show a slightly smaller NE population at 48 h AEL, likely indicating that retarded NE proliferation begins earlier. By comparison, the NE population of *A-indel* mutants looks normal at 48 h AEL but shows retarded growth at 72 h AEL, implying that BaboA function becomes necessary in late period of the first phase. Although it is not known if the first phase of NE expansion can be further divided into two distinct periods, our observations imply that BaboA might function differently in NEs during early and late periods of the first phase, while pan-Babo function is essential throughout. In this aspect, it is interesting to note that EcR-B1 expression is detected first around 66 h AEL (L2) in the OL NEs [32]. The similar timing of BaboA’s requirement during the late first phase proliferation of NEs and on EcR-B1 expression in the OL NEs might not be coincidental but instead might support a causative relationship between the two.

The OPC NEs in the second phase transform into the medulla NBs medially and the LPCs laterally [24]. However, both pan-*babo* and *A-indel* mutants contain a small number of medulla NBs and LPCs at 120 h AEL, suggesting a conversion defect of mutant OPC NEs. Cell- and stage-specific knockdown assays of *baboA* demonstrated an autonomous role of BaboA for this cellular event in the OPC. Unlike OPC-NEs, IPC-NEs first transform into migratory progenitor cells that travel distally in streams and then turn into NBs upon arrival at their destination (d-IPC zone), and ultimately give rise to the lobula plate [50]. Knockdown of *baboA* in IPC suppressed growth in the d-IPC zone, suggesting that BaboA also plays an autonomous role in the conversion of IPC NEs. Despite BaboA’s essential role in NE conversion, the more severe defect of pan-*babo* mutants in this cellular event again implies actions of other isoforms. Currently, the molecular mechanisms underlying the conversion of IPC NEs are poorly understood but our studies indicate that IPC NEs also require Babo function in a similar manner as OPC NEs.

### Babo isoforms regulate stage-specific neural expression of EcR-B1

Ecdysone/EcR signaling is important for the initiation of OPC NE conversion and CB-NBs entering the second phase proliferation mode [7]. Since *babo* is a key factor regulating the stage-specific expression of EcR-B1 in the larval CNS [18], it is conceivable that *babo* promotes second-phase behaviors of CB-NBs and OL-NEs through EcR-B1 expression. In support of this, EcR-B1 expression levels are greatly reduced in the CNS of *baboA* mutants but clearly to a lesser degree than pan-*babo* mutants. It appears that BaboA promotes stage-specific upregulation of neural EcR-B1 expression to some extent, while combined actions of multiple isoforms are required to raise the neural EcR-B1 expression to a peak level. In support, blocking of EcR function in both CB-NBs and OL-NEs causes severe reduction in both brain regions, which are much more severe than *baboA*-KD. Taking these observations into consideration, we propose that Babo isoforms act together to upregulate stage-specific expression levels of EcR-B1 in both neural stem cells, which in turn transmits the ecdysone signaling to promote the second phase behavior of both types of neural stem cells.

## Materials and Methods

### Fly strains

Fly lines were raised on standard cornmeal-yeast-agar medium at 25°C. A *trans*-heterozygous combination of *babo^Fd4^* and *babo^Df^* (a.k.a. *bobo^NP4^* [34], *bobo^β^* [39]) was used as a pan-*babo* mutation. The following mutant alleles were also used: *myo^β1^* [41], *myo^CR2^* [56], *dSmad2^F4^* (a.k.a. *Smox^F4^*) [44]. All mutant alleles were balanced with either *act-GFP* or *unc13-GFP*. *w^1118^* strain and *babo^Df/+^* were used as wild-type control and genetic control, respectively. The latter was obtained by crossing *babo^Df^/CyO act-GFP* with *w^1118^*. For transgenic manipulations, *UAS-baboA-miRNA* [18], *UAS-EcR-B1^W650A^*(*EcR-DN*) [57], *daughterless (da)-gal4* [58], *R31H09-gal4* (BL-49694), *UAS-dSmad2-RNAi* (BL-41670), *scarecrow* (*scro*)-*gal4* [52], and *tub-gal80^ts^* (BL-7017) were used. BL indicates the stocks obtained from the Bloomington Drosophila Stock Center (https://flystocks.bio.indiana.edu/).

### Generation of transgenic lines

To generate two independent deletion mutations for each *babo* isoform, a CRISPR mutagenesis approach was employed to delete target sequences within isoform-specific 4^th^ exons. The DNA lesions for the six alleles are shown (S1 Fig). Primer pairs (S1 Table) were selected using the CRISPR Optimal Target Finder tool and cloned into pU6-BbsI [59]. Plasmids expressing gRNA were injected into a Cas9-expressing stock (GenetiVision; www.genetivision.com/). PCR products containing the gRNA target site were generated from heterozygous males and the molecular lesions were identified by comparing the observed mixed sequence to the wild-type sequence of the balancer.

### Growth of mutant larvae

Flies were allowed to lay eggs for 2-3 h on an apple juice plate supplemented with yeast paste, and then the embryos were raised in a humidified chamber at 25°C under 12 h light/12 h dark cycles. About 35 homozygous mutant larvae lacking balancer-derived GFP expression were collected soon after hatching and raised as a group in a plate containing low sugar apple juice medium [1X apple juice (Old Orchard), 2.5% glucose, 2.2% agar, and 0.15% of methyl paraben] with extra yeast paste to avoid overcrowding and sugar toxicity that is detrimental to pan-*babo* mutants [40].

### Immunohistochemistry and imaging

Immunohistochemical procedures followed essentially the same method as described [60], except for the treatment with primary antibodies for 2 h at room temperature (RT) with shaking. DAPI staining was done by treating the tissues in a dilution of 1:500 for 40 min. The processed CNS tissues were mounted with a shielding medium (2% n-propyl galate in 80% glycerol). The following primary antibodies were used; rabbit anti-GFP (1:500) [61], mouse anti-24B10 (1:100, Developmental Studies Hybridoma, Bank, DSHB), mouse anti-Dlg (1:100, DSHB 4F3), rat anti-Mira (1:100, Abcam), mouse anti-Dac (1:100, DSHB mAbdac2-3), and anti-EcR-B1 (1:50, DSHB AD4.4). Secondary antibodies conjugated with Alex Fluor 594 (1:200; Jackson ImmunoResearch Lab), Alex Fluor 555 (1:200, Molecular Probes), or Alex Fluor 455 (1:200, Molecular Probes) were used. Confocal images were obtained with a Zeiss LSM710 or Leica SP8 confocal microscope and epi-fluorescent images with an Olympus BX61 microscope. Fly body images were taken with a Zeiss Semi stereo microscope. Images and figures were generated by using ImageJ (NIH) and Microsoft PowerPoint, respectively.

### Measuring areas of brain regions and body lengths

Labeled larval brains were mounted dorsal between 1-mm-thick 18X18mm coverslips (spacers) on a slide glass, covered with another cover slip on top, and sealed with clear nail polish. A frontal mid-section (2 μm thick) image that was taken from a lobe per brain. Areas of different brain regions were manually measured on individual frontal mid-section images using the ImageJ program. To measure larval body length, the whole-body images were taken under same condition after killing larvae by dipping into in boiling water for a few seconds. All measurements in graphs are in pixels or as otherwise indicated.

### Statistical analysis and graphing

Statistical analyses were performed using GraphPad Prism (9.0v). Ordinary one-way ANOVA followed by Tukey’s multiple comparisons was used for more than three groups and unpaired student t-test for comparison between two groups. Different numbers of asterisks denote statistical significance with the following p-values: *p<0.05, ** p<0.01, ***p<0.001, ****p<0.0001, and ns for not significant (p>0.05). Graphs were generated by using either GraphPad Prism or Microsoft Excel. All bars in graphs show mean ± sd (standard deviation).

## Acknowledgments

We greatly appreciate the efforts of Dr. Shimell for comments on the manuscript, Dr.Bretscher for technical advices, and David Zhitomirsky for help with molecular cloning. We also thank Dr. Goodchild for the kind gift of the anti-GFP.

## Author Contributions

**Conceptualization:** Gyunghee G. Lee, Aidan J. Peterson

**Formal analysis:** Gyunghee G. Lee

**Funding acquisition:** Jae H. Park, Michael B. O’Connor

**Investigation:** Gyunghee G. Lee, Aidan J. Peterson, Myung-Jun Kim

**Project administration:** Jae H. Park, Michael B. O’Connor

**Resources:** Jae H. Park, Michael B. O’Connor

**Supervision:** Jae H. Park, Michael B. O’Connor

**Visualization:** Gyunghee G. Lee, Aidan J. Peterson, Myung-Jun Kim

**Writing – original draft:** Gyunghee G.Lee

**Writing – review & editing:** Gyunghee G. Lee, Aidan J. Peterson, Jae H. Park, Michael B. O’Connor

## Supporting information

**S1 Table.** Primer sequences used for making guide RNA constructs targeting isoform-specific 4^th^ exons and for PCR confirmation.

**S1 Fig. Deduced sequences of CRISPR-generated isoform-specific mutant alleles.**

Wild-type (WT) sequences near the respective guide sites are shown for comparison. First nucleotide shown is G1168 for *babo*-A (NM_057652.4), C1159 for *babo*-B (NM_057651.4), and T1141 for *babo*-C (NM_001169621.2). Each allele lists the name used in the main text, the original chromosome name, the sequencing reaction used to read the altered sequences, and a description of the resulting changes. f1, f2, of f3 refers to the reading frame following the lesion. ^ indicates one or more coding triplets not normally found in the gene. All lesions are simple deletions except: *A-indel* (*AΔ9*) results from Δ12 with insertion of 3 extra nucleotides, and *C-indel* (*CΔ1*) results from Δ3 with insertion of 2 extra nucleotides.

**S2 Fig. NB populations in the central brain region of *pan-babo* and *A-indel* mutants.** The brains at 120 h AEL were simultaneously processed for anti-Mira immunostaining (n=7 for each genotype). About 3-4 optical Z-sections (2 μm/each) were made into a composite image to view a population of NBs in ventral **(A-C)** and dorsal side **(Ai-Ci)** of each brain lobe. Scale bar in A, 50 μm.

**S3 Fig. *baboA-KD* by *scro-gal4* in subpopulation of medulla NBs did not cause any significant proliferation defect.** *UAS-mCD8GFP; scro-gal4* flies were crossed to *w^1118^*(con) or *UAS-babo-miRNA* for *baboA*-KD. F1 brains were examined for GFP expression in the ventral side of the OL at WL3. Strong GFP expression driven by *scro-gal4* is detected in the subpopulation of medulla NBs and continuously in their clones (brackets) and lamina neurons but not in any of OPC NEs. **(A)** con (n=5). **(B)** *baboA-KD* (n=5). Scale bar in A: 50 μm.

**S4 Fig. Reduction in medulla NB population due to *baboA*-KD and overexpression of EcR-DN in NEs at WL3 stage.** The medulla fields are indicated by asterisks and the centers of distal-IPC zones by red arrowheads. **(A)** *R31H09*>mCD8*GFP(GFP)*+*nRFP* (con, n=13). *R31H09-gal4* drove GFP expression strongly in NEs but little or none in other cell types of the OL, while nRFP expression was detected strongly in NEs and weakly in the precursor cells. Small strips of OPC and IPC NE layers are in stark contrast with large medulla field and d-IPC zone. **(B)** *R31H09*>*mCD8GFP*+*nRFP*+*mibaboA* (*baboA*-KD, n=15). **(C)** *R31H09*> *mCD8GFP*+*nRFP*+*EcR-DN* (EcR-DN, n=12). **(D-F)** Mira positive medulla NBs on the ventral side of the brain lobe are indicated by brackets. **(G)** Mean areas (± sd) of the brain lobe (black), CB (blue), and OL (red) regions. Scale bars in A and in D: 50 μm.

